# Multicomponent *Pseudomonas aeruginosa* vaccines eliciting Th17 cells and functional antibody responses confer enhanced protection against experimental acute pneumonia in mice

**DOI:** 10.1101/2022.05.05.490820

**Authors:** Mohammad Omar Faruk Shaikh, Matthew M. Schaefers, Christina Merakou, Marco DiBlasi, Sarah Bonney, Tiffany Liao, David Zurakowski, Margaret Kehl, David E. Tabor, Antonio DiGiandomenico, Gregory P. Priebe

**Author notes:** Mohammad Omar Faruk Shaikh and Matthew M. Schaefers contributed equally to this work. Author order was determined in order of increasing seniority. Corresponding authors Mailing address: Division of Critical Care Medicine, Department of Anesthesiology, Critical Care and Pain Medicine, Boston Children’s Hospital, 300 Longwood Ave., Boston, MA 02115, USA., and. Italian National Health Institute (ISS), Rome, Italy.

## Abstract

The Gram-negative pathogen *Pseudomonas aeruginosa* is a common cause of pneumonia in hospitalized patients. Its increasing antibiotic resistance and widespread occurrence present a pressing need for vaccines. We previously showed that a *P. aeruginosa* type III secretion system protein, PopB elicits a strong Th17 response in mice after intranasal (IN) immunization and confers antibody-independent protection against pneumonia in mice. In the current study, we evaluated the immunogenicity and protective efficacy in mice of the combination of PopB (purified with its chaperone protein PcrH) and OprF/I, an outer membrane hybrid fusion protein, compared to immunization with the proteins individually either by the intranasal (IN) or subcutaneous (SC) routes. Our results show that after vaccination, a Th17 recall response from splenocytes was detected only in mice vaccinated with PopB/PcrH, either alone or in combination with OprF/I. Mice that were immunized with the combination of PopB/PcrH and OprF/I had enhanced protection in an acute lethal *P. aeruginosa* pneumonia model, regardless of vaccine route, compared to the mice vaccinated the with either alone or adjuvant control. Immunization generated IgG titers against the vaccine proteins and whole *P. aeruginosa* cells. Interestingly, none of these antisera had opsonophagocytic killing activity, but antisera from mice immunized with vaccines containing OprF/I had the ability to block IFN-γ binding to OprF/I, a known virulence mechanism. Hence, vaccines combining PopB/PcrH with OprF/I that elicit functional antibodies lead to a broadly and potently protective vaccine against *P. aeruginosa* pulmonary infections.

## Introduction

The Gram-negative bacterium *Pseudomonas aeruginosa* causes a wide range of clinically important infections, mostly in hospitalized and immunocompromised patients, especially those requiring mechanical ventilation, those with burns or combat-related wounds, and in people with cystic fibrosis (CF). *P. aeruginosa* is the most common pathogen causing ventilator-associated pneumonia (VAP) worldwide, with a prevalence of 3–5% in adults ventilated for more than 48 hours (1). The need for an effective vaccine for *P. aeruginosa* VAP is made even more urgent due to COVID-19, where the rate of VAP is strikingly high (2, 3). The steadily increasing antibiotic resistance encountered in *P. aeruginosa* clinical isolates (4-6), coupled with the relative dearth of new antibiotics in the pharmaceutical industry’s pipeline, also make paramount the need for new approaches to the development of an effective vaccine.

Although antibodies to the lipopolysaccharide (LPS) O antigen mediate high-level immunity to *P. aeruginosa* infections, vaccine strategies targeting the O antigen have not been successful to date and are stymied by O antigen variability and by impaired immune responses when O antigens of different serotypes are combined (7-12). More recent *P. aeruginosa* vaccine strategies showing promise have focused on the outer membrane proteins OprF, OprI, and OprF/I fusion proteins (specifically OprF_190-342_-OprI_21-83_ called VC43, previously IC43), with and without flagellins (13-18). The OprF/I fusion protein vaccine VC43 remains a promising candidate as it induces multiple immune effectors in mice, including opsonic antibodies (18), antibodies that inhibit IFN-γ binding to *P. aeruginosa* (19) (thereby interfering with a virulence mechanism (20)), and, via the OprF_329-342_ epitope, IFN-γ^+^ T cell responses (15). However, despite positive results in a phase II trial demonstrating improved all-cause mortality (21), in a phase III trial targeting VAP prevention, VLA43 failed (22). We speculate that the failure of the VLA43 trial was related to the absence of an adjuvant and/or the lack of induction of a Th17 response, which our published (23, 24) studies show is critical for vaccine-induced broad protection.

Our previous work has identified PopB, a structural component of the type III secretion system (T3SS), as a protective T cell antigen that generates a Th17 response when administered intranasally with the Th17 adjuvant curdlan (23, 24). The *popB* gene has been found in nearly all *P. aeruginosa* strains (25, 26), and PopB is highly conserved and expressed during infection (25, 27, 28). A recombinant His-tagged version of PopB is soluble and stable only when co-purified with its chaperone PcrH, so PopB is denoted PopB/PcrH. Our previous work has found that immunization with PcrH does not elicit a Th17 response and the addition of PopB is required for full protection (23, 24).

Here we have studied the immune responses and protective efficacy against pneumonia in mice after vaccination with PopB/PcrH, OprF/I, or an admixture of both, via the intranasal (IN) or subcutaneous (SC) immunization routes. We report that the combined vaccine yields the highest protection regardless of route, likely due to eliciting both Th17 and functional antibody responses, particularly antibodies that inhibit the binding of OprF to IFN-γ.

## Results

### Immunization of mice with PopB-containing vaccines elicits a Th17 response, while OprF/I does not

With the failure of the phase III clinical trial using OprF/I as the vaccine antigen, novel approaches to vaccinate against *P. aeruginosa* infections are needed. Our previous work demonstrate that an effective Th17 response is required for robust protection against *P. aeruginosa* infections (23). We hypothesized that the OprF/I vaccine used in the recent clinical trial was unable to induce a Th17 response, and potentially why it failed. To test this hypothesis, mice were immunized with PopB/PcrH, OprF/I, a combination of PopB/PcrH and OprF/I, or adjuvant alone. Our previous work with PopB has found that its chaperon, PcrH, is needed for effective expression and purification. Both proteins PopB and PcrH are used in immunization, but only PopB generates a Th17 response, and that response is protective (23, 24). We prepared the recombinant fusion protein consisting of OprF_190-342_ and OprI_21-83_ as described for the OprF/I vaccine VLA43 (18). Mice were immunized IN using the Th17 adjuvant curdlan (29) or SC using aluminum hydroxide (Alum) as adjuvant. Three weeks after the last immunization, splenocytes were isolated and stimulated with either PopB/PcrH or OprF/I, and IL-17 was measured to quantify the Th17 response elicited by each vaccine. Splenocytes from mice immunized with OprF/I did not produce IL-17 when stimulated with either OprF/I or PopB/PcrH when immunized via an IN route (Figure 1A) or a SC route (Figure 1B) demonstrating that OprF/I does not induce a Th17 response in this mouse strain. Splenocytes that were isolated from mice that were immunized with PopB/PcrH, either alone or in combination with OprF/I produced IL-17 when stimulated with PopB/PcrH (Figure 1). There were statistically significant (*p* value <0.05) increases in amounts of IL-17 produced in splenocytes isolated from mice immunized with the combination of PopB/PcrH and OprF/I compared to mice immunized with PopB/PcrH alone when immunized via IN or SC routes (∼1.6-fold and ∼5-fold, respectively, Figure 1) suggesting no evidence of inhibition or anergy when the OprF/I was admixed with PopB/PcrH.

**Figure 1.**
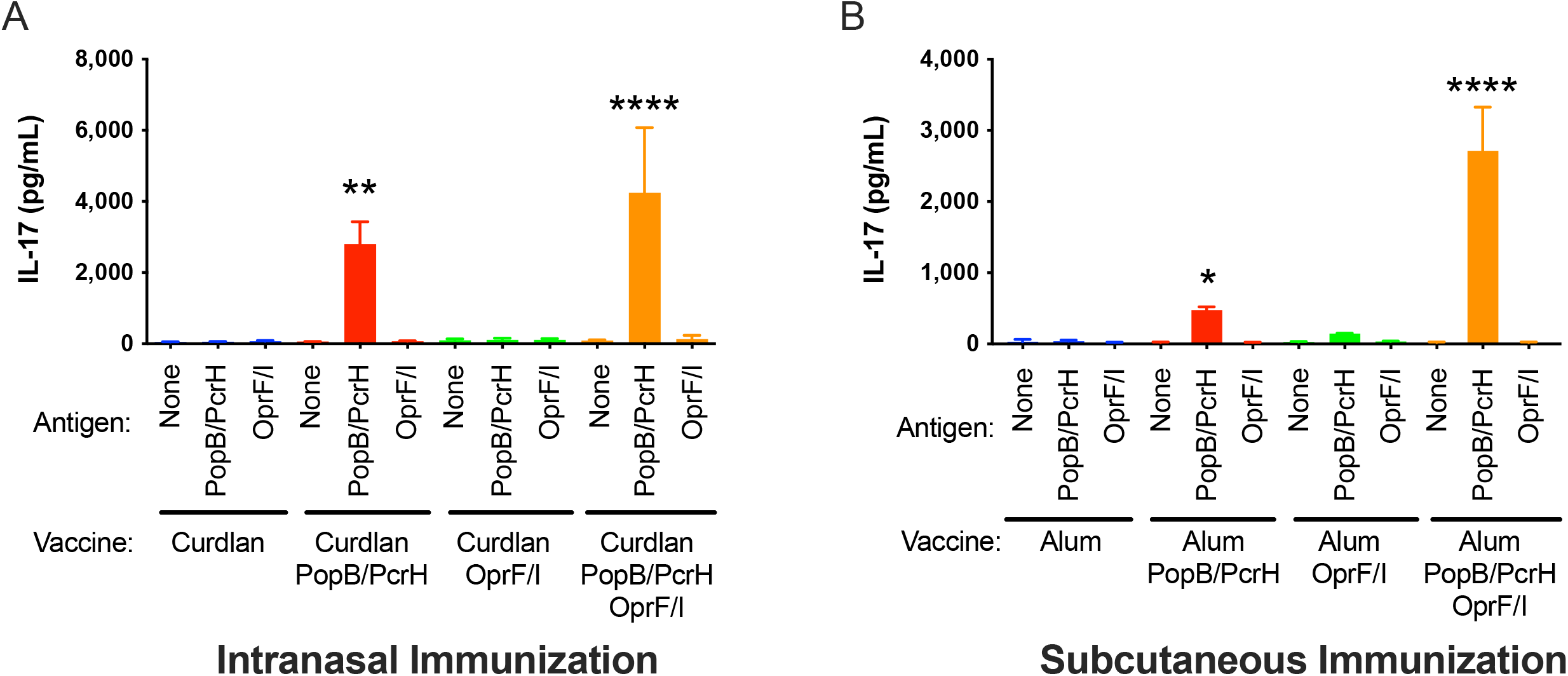
Vaccination of mice with PopB/PcrH but not OprF/I generates a Th17-response. Splenocytes from mice that were immunized either intranasally (A) or subcutaneously (B) with adjuvant alone (curdlan or Alum), adjuvant + 30 µg PopB/PcrH, adjuvant + 30 µg OprF/I, or adjuvant + 30 µg PopB/PcrH + 30 µg OprF/I were isolated 3 weeks after last immunization and stimulated with either PopB/PcrH, OprF/I or left unstimulated. IL-17 was measured by ELISA after 7 days of stimulation. Bars are the average of triplicates using pooled spleens from 4 mice, and error bars are SDs, and are representative of experiments conducted at least two times. * *p*<0.05, ** *p*<0.01,*** *p*<0.001, \*\*\*\**p*<0.0001 by one-way ANOVA followed by Dunnett’s multiple comparison test when compared to the adjuvant only vaccine group (curdlan or Alum) stimulated with media.

### Immunization with the combination of PopB/PcrH and OprF/I induces significant protection against acute *P. aeruginosa* infection

We next evaluated the protection provided by PopB/PcrH and OprF/I-based vaccines against acute *P. aeruginosa* pneumonia after intranasal inoculation. Three weeks after the final immunization mice were challenged with a lethal dose (>LD_100_) of *P. aeruginosa* strain N13 (2×10^6^ CFU/mouse), and the survival was monitored for 6 days. Mice that were immunized with the combination of PopB/PcrH and OprF/I showed significant protection against challenge compared to adjuvant alone in both the IN and SC routes (*p* values = 0.014 and 0.008, respectively, by log-rank test, Figure 2). Mice that were immunized either IN or SC with PopB/PcrH or OprF/I alone were not significantly protected compared to adjuvant alone, demonstrating the combination of vaccine antigens conferred greater protection than either of the antigens alone. It is worth pointing out a higher dose of a different *P. aeruginosa* strain was used in our current study compared to our previously published findings using PopB/PcrH vaccines, which used the challenge strain ExoU^+^ PAO1, a highly virulent engineered strain expressing the ExoU cytotoxin and its chaperone by a plasmid (23, 24).

**Figure 2.**
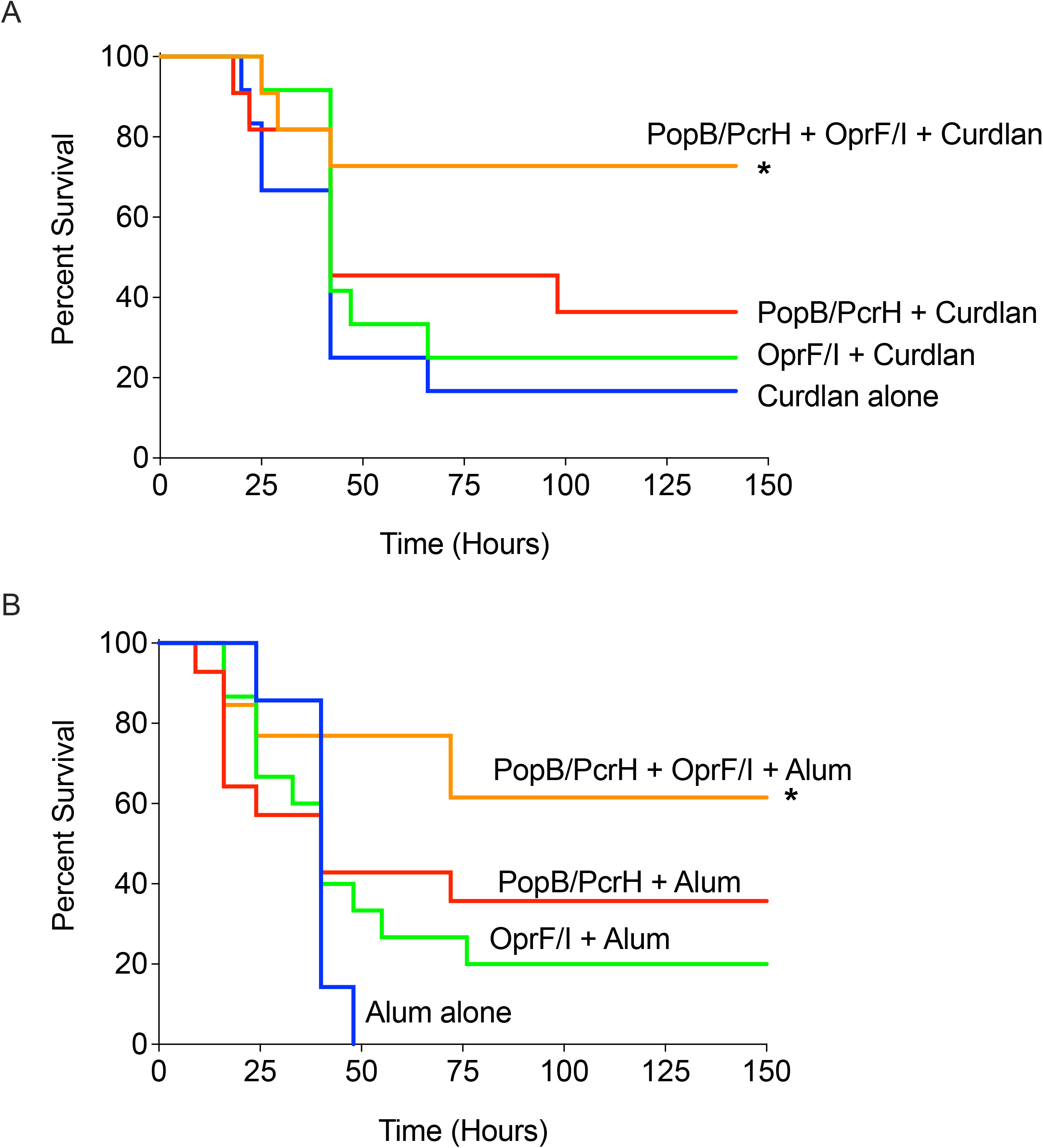
The combination of PopB/PcrH and OprF/I protects against acute lethal *P. aeruginosa* pneumonia. Mice that were immunized either intranasally (A) or subcutaneously (B) with adjuvant alone (curdlan or Alum), adjuvant + 30 µg PopB/PcrH, adjuvant + 30 µg OprF/I, or adjuvant + 30 µg PopB/PcrH + 30 µg OprF/I were challenged with *P. aeruginosa* strain N13 (2×10^6^ CFU/mouse) three weeks after the last immunization. Disease progression was monitored over 6 days, * denotes *p* value <0.05 by log-rank test compared to adjuvant alone. n=11-12 mice per group, data are pooled from at least two independent experiments.

### Intranasal and subcutaneous immunization with PopB/PcrH or OprF/I elicit IgG responses

To better describe the immune response generated by the various vaccines, we measured the humoral immune responses after immunization. Sera from immunized mice were collected 3 weeks after the third immunization, and IgG titers specific for PopB, OprF/I, and whole *P. aeruginosa* were measured (Figure 3 and Table S1A). As expected, mice generated IgG responses against the proteins they were immunized against. In IN-immunized mice, there was no difference in the EC_50_s of anti-PopB IgG titers in mice that were immunized with PopB/PcrH alone compared to combination of PopB/PcrH and OprF/I, when comparing the 95% confidence intervals of EC_50_ determinations. Likewise, there was no difference in the anti-OprF/I titers in mice that were immunized with OprF/I alone compared to combination of PopB/PcrH and OprF/I. Among SC-immunized mice, there was significant 1.7-fold increase in EC_50_s of anti-OprF/I IgG titers in mice that were immunized with the combination of PopB/PcrH and OprF/I compared to mice immunized with OprF/I alone. Conversely, the anti-PopB titers were 10-fold lower in mice that were immunized with the combination of PopB/PcrH and OprF/I compared to the PopB/PcrH only group. We also measured the IgG titers against whole *P. aeruginosa* bacterial cells, using strain N13, which was the same strain that was used in challenge experiments. In mice that were IN immunized, there was a significant ∼2 or 2.7-fold increase in the EC_50_s of anti-*P. aeruginosa* IgG titers in mice that were immunized with the combination of PopB/PcrH and OprF/I compared to mice that were immunized with PopB/PcrH or OprF/I alone, respectively. In the SC-immunized mice, immunization with the combination of PopB/PcrH and OprF/I or OprF/I alone resulted in a ∼200-fold significant increase of anti-*P. aeruginosa* IgG titers than the mice that were immunized with PopB/PcrH alone. We also measured the anti-PopB or anti-OprF/I titers of the IgG1 and IgG2c subclasses. In mice, a strong IgG1 response is associated with a Th2 response while a IgG2c response is associated with a Th1 response (30). Mice that were immunized via the SC route had higher IgG1 titers to both PopB and OprF/I suggesting a Th2 response (Figure S1 and Table S1B).

**Figure 3.**
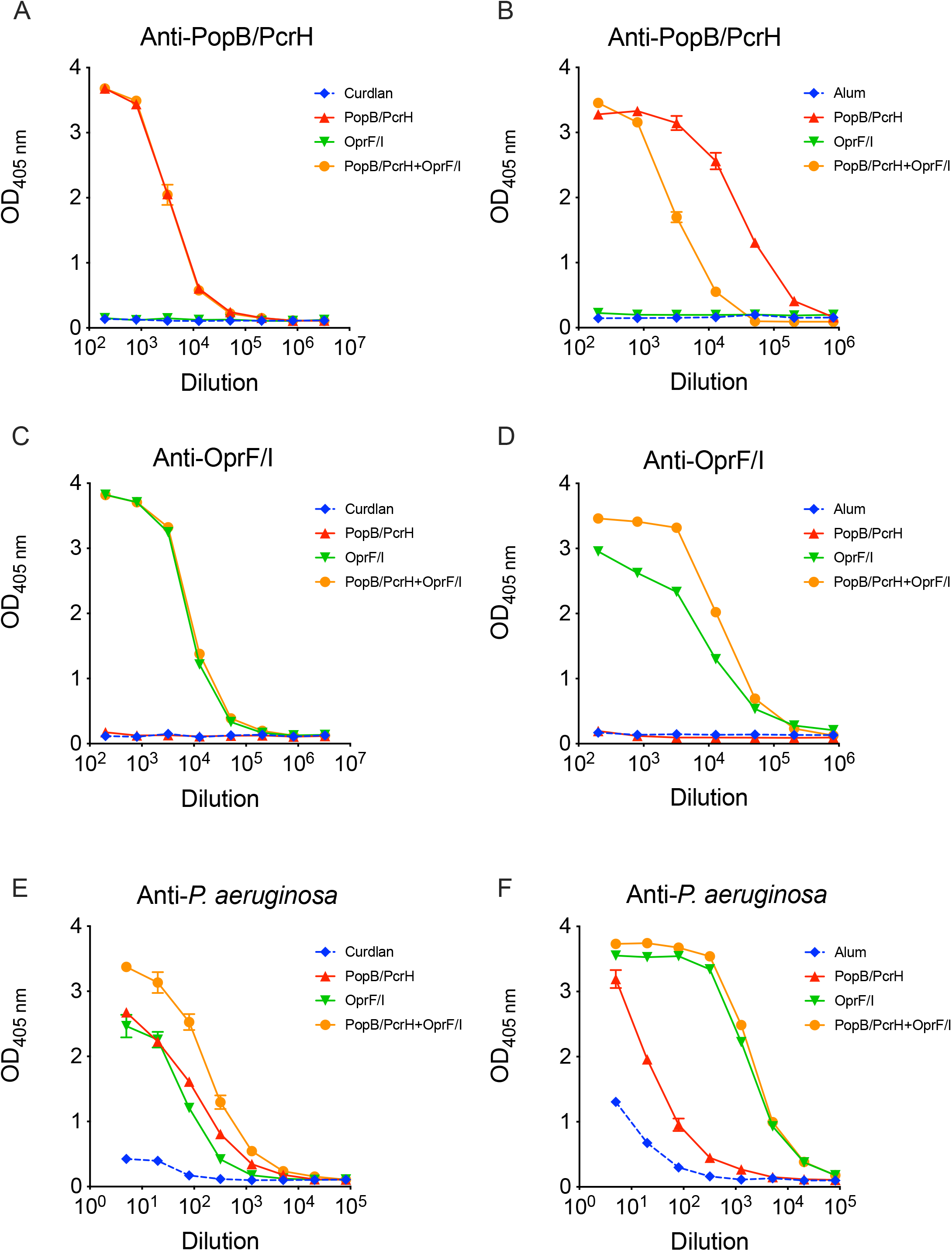
Vaccination with PopB/PcrH, OprF/I, or both, generate antigen-specific IgG responses that also recognize whole *P. aeruginosa*. Mice were immunized either intranasally (A,C,E) or subcutaneously (B,D,F) with adjuvant alone (curdlan or Alum), adjuvant + 30 µg PopB/PcrH, adjuvant + 30 µg OprF/I, or adjuvant + 30 µg PopB/PcrH + 30 µg OprF/I and sera were collected three weeks after the last immunization. Anti-PopB/PcrH (A,B), anti-OprF/I (C,D), and anti-whole *P. aeruginosa* strain N13 (E,F) IgG titers were measured using ELISA. Sera from 3-4 mice per group were pooled and measured in duplicate, and means are plotted with SD as error bars (error bars are smaller than symbol at many points). Data are representative of at least two independent experiments.

### Sera from mice immunized with OprF/I inhibit OprF binding to IFN-γ but lack opsonophagocytic killing (OPK) activity

*P. aeruginosa* OprF can bind to human IFN-γ thereby enhancing virulence (20). Furthermore, Anti-OprF/I antibodies are able to prevent the fusion protein OprF/I from binding to IFN-γ *in vitro*, which has been suggested as one mechanism that vaccination with OprF/I can prevent infection (19). Thus, we measured the ability of various sera to block OprF/I binding to human IFN-γ. Sera from mice that were immunized with OprF/I either alone or with PopB/PcrH had a statistically significant 25-33% reduction in OprF/I binding to IFN-γ when vaccinated either IN or SC (Figure 4). Sera from mice that were immunized with adjuvant or PopB/PcrH alone did not inhibit OprF/I binding to IFN-γ. Surprisingly sera from OprF/I-immunized mice did not have OPK activity (Figure S2) against target strain PAO1 (serotype O2/O5) and strain 9882-80 (serotype O11), perhaps due to using a different mouse strain compared to prior published work (31), Our previous results have shown that anti-PopB/PcrH antibodies do not mediate OPK (23, 24), and the current study confirmed those results (Figure S2).

**Figure 4.**
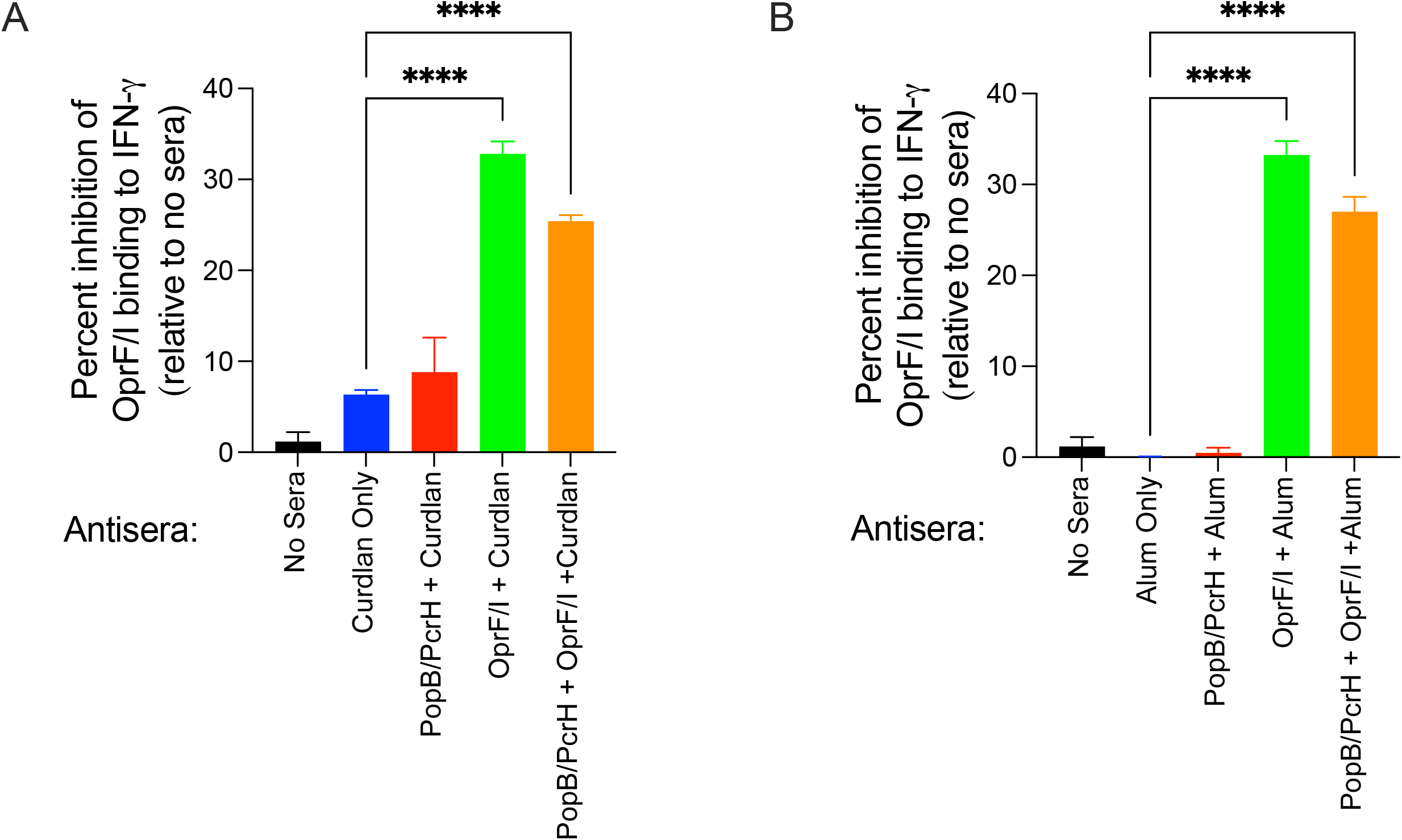
Sera from mice immunized with OprF/I inhibit binding of OprF/I to human IFN-γ. Sera were collected three weeks after the final immunization of mice that were immunized either intranasally (A) or subcutaneously (B) with adjuvant alone (curdlan or Alum), adjuvant + 30 µg PopB/PcrH, adjuvant + 30 µg OprF/I, or adjuvant + 30 µg PopB/PcrH + 30 µg OprF/I. Pooled sera were diluted 1:2 and assessed for inhibition of binding of human IFN-γ to OprF/I coated on wells of ELISA plates. IFN-γ bound to OprF/I was measured using anti-IFN-γ antibodies. The percent inhibition was calculated as the change in the amount of IFN-γ that bound OprF/I in the absence of sera. Sera from 3-4 mice per group were pooled and measured in triplicate, and means are plotted with SD as error bars. ****denotes *p* value <0.0001 by one-way ANOVA followed by the Sidak multiple comparison test. Data are representative of at least two independent experiments.

## Discussion and Conclusions

An effective vaccine to prevent *P. aeruginosa* infections remains an unrealized goal that could prevent a significant amount of morbidity and mortality (8, 12). The failure of recent clinical trials highlights the need for novel approaches. Our previous work demonstrated that IL-17 is critical in protection against LPS-heterologous strains of *P. aeruginosa* after vaccination with live-attenuated *P. aeruginosa* vaccines (31) and identified PopB as a protein stimulating Th17 responses and as promising vaccine candidate for *P. aeruginosa* (23). In this study we generated a novel vaccine that combined PopB/PcrH along with a recombinant fusion protein consisting of portions of OprF and OprI and evaluated vaccination via the IN and SC routes in mice using a combination of the antigens as well as the individual antigens.

We found that immunization with PopB-containing vaccines (either IN or SC) resulted in a Th17 response (Figure 1). While OprF/I alone did not induce a Th17 response there was significant increase in the Th17 response when OprF/I was added to PopB/PcrH suggesting an adjuvant-like effect of unclear mechanism. These Th17-responses are consistent with findings that both we and other groups have observed when immunizing with PopB-containing vaccines (23, 24, 32). Th17 responses are essential for host defense against a number of pathogens, including *Salmonella enterica* (33), *Streptococcus pneumoniae* (34), *Klebsiella pneumoniae* (35), *Staphylococcus aureus* (36, 37), *Mycobacterium tuberculosis* (38), and *Candida albicans* (39).

Vaccination with the combination of PopB/PcrH along with OprF/I resulted in significant protection against a lethal challenge of *P. aeruginosa* in our murine pneumonia model (Figure 2). In the current study we used a higher dose of *P. aeruginosa* strain N13, a clinical isolate of serotype O6) compared to our previous studies where we used *P. aeruginosa* strain PAO1 (serotype O2/O5) that expresses ExoU from a plasmid (23, 24). Immunization with either PopB/PcrH or OprF/I alone was not sufficient to protect mice against the lethal challenge doses used in this study; but the combination vaccine protected mice, whether vaccinated IN or SC. Our previous work has identified that the Th17 response is required for broad serotype-heterologous protection against *P. aeruginosa* infection (23), and such Th17 responses are likely contributing to the protection observed in the current study.

Immunization with PopB/PcrH, OprF/I, or the combination of the two proteins elicited antigen-specific IgG antibody responses. The immunizations also elicited an antibody response that was able to recognize whole *P. aeruginosa* coated on ELISA plates. The contributions to protection of these antibodies remains unclear. Our previous work, and current work with PopB-based immunization, demonstrate that antibodies generated in response to PopB do not have OPK activity (Figure S2) (23, 24). Studies in humans (18), nonhuman primates (40), and other mouse strains (14) found that immunization with OprF/I generates antibodies with OPK activity, but we observed no such activity in the current study. The overall response to the OprF/I was IgG1 biased suggesting a Th2 response, which is also consistent with a previous study (41). We did find that mice immunized with OprF/I generated antibodies that were able to block the binding of OprF/I to IFN-γ, a host-sensing mechanism shown to enhance the virulence of *P. aeruginosa* (20). These antibodies that prevent the OprF-IFN-γ interaction are predicted to be one mechanism that immunization with OprF/I can reduce *P. aeruginosa* virulence (19).

Based on the failure of the OprF/I vaccine (VLA43) in phase III clinical trials, an effective vaccine that can prevent *P. aeruginosa* infections will need to elicit multiple mechanism of action, with Th17 responses as a critical component. The current work suggests that combining PopB with OprF/I significantly improves protective efficacy against acute lethal pneumonia in mice compared with either protein alone, advancing us one step closer to a broadly and potently protective vaccine for *P. aeruginosa*.

## Materials and Methods

### Generation of protein expression vectors

A gene fragment encoding *E. coli-*codon-optimized OprF_190-342_ and OprI_21-83_ was synthesized with a 5 prime *ndeI* and a 3 prime *xhoI* restriction site (Genescript). This fragment was then cloned into the *ndeI* and *xhoI* site of pET24a(+) creating pET24a(+)-OPRF/I. Insert was confirmed by Sanger sequencing. The pET28b-based PopB/PcrH expressing vector was previously described (23).

### Expression and purification of OprF/I and PopB/PcrH from *E. coli*

Purification of OprF/I was performed as previously described in patent filings (https://patents.google.com/patent/EP2686339A1/). Briefly, *E. coli* BL21(DE3) carrying pET24a(+)-OPRF/I was grown in 2L of fresh LB media with kanamycin and grown at 37C. When an OD_600_ of 0.8 was reached, the culture was induced with 1 mM of IPTG (isopropyl _β_-d-1-thiogalactopyranoside) and incubated for an additional 3.5 hours. *E. coli* cells were then harvested using centrifugation for 10 minutes at 7800g in 4C. Cells were resuspended in lysis buffer (Buffer A) containing 8M urea, 20 mM Tris-HCl, 100 mM KCl, 200 mM NaCl, and 10 mM Imidazole and then sonicated. Lysates of cells were centrifuged at 10,000g for 30 minutes and supernatants were purified through an IMAC column. Column resin was washed with 50 column volumes of Buffer A containing 0.1% Triton X-114 followed by 20 column volumes of Buffer A. OprF/I was then eluted in Buffer A containing 250 mM Imidazole. Fractions of elution underwent refolding dialysis with urea, endotoxin removal with polylysine resin, and then final dialysis with reoxidation of the purified material using 1 mM DTT. PopB/PcrH was purified as previously described (23).

### Immunization of Mice and Murine Pneumonia Model

All animal protocols and procedures were approved by the Boston Children’s Hospital Institutional Animal Care and Use Committee (assurance number A3303-01). The specific protocol numbers are 18-01-3617R and 20-12-4326R. All animal protocols are compliant with NIH Office of Laboratory Animal Welfare, Guide for the Care and Use of Laboratory Animals, The US Animal Welfare Act, and PHS Policy on Humane Care and Use of Laboratory Animals. FVB/N mice (6–8 weeks old, female) were obtained from the Jackson Laboratories and maintained in house for both vaccination periods and challenge experiments.

Vaccine formulation and immunization: SC formulated vaccines contained in 200 µL per dose of the following: 30 µg of each protein (PopB/PcrH, OprF/I, or both) mixed with Alum (Alhydrogel®; aluminum hydroxide adjuvant 2%, 25 µL, 250 µg/dose; InvivoGen) and suspended in saline with incubated for 1 hour at room temperature with gentle rotation for adsorption prior to vaccination. SC vaccination was conducted once every two weeks for a total of 3 vaccinations (Day 0, 14, and 28) with Alum alone as negative control (30). Vaccines formulated for IN immunization contained in 30 µL per dose (15 µL per nostril) the following: 30 µg of each protein (PopB/PcrH, OprF/I, or both) mixed with curdlan (150 µg/dose) (Sigma-Aldrich) and suspended in saline. Vaccination was conducted once every week for a total of 3 vaccinations (Day 0, 7, and 14) with curdlan alone as negative control (31).

### Mouse sera collection

Blood samples from mice were collected retro-orbitally three weeks after the 3^rd^ vaccination [i.e. Day 35 (for IN) and Day 56 (for SC), where Day 0 is the 1^st^ vaccine dose]. The sera were separated using serum separator tubes (BD). Sera were aliquoted and stored at -80°C until use.

### Challenge experiments

Mice were challenged with *P. aeruginosa* using previous published methods (23, 24). Briefly, *P. aeruginosa* strain N13 was grown overnight from a frozen stock at 37°C on tryptic soy agar (TSA) plates. Bacteria scraped from the plate were suspended in 10 mL PBS (Invitrogen) to OD_650nm_ of 0.55, which corresponds to approximately 10^9^ CFU/mL. Bacterial suspensions were diluted to prepare the inocula, and bacterial doses were confirmed by serial dilution and plating. Mice were anesthetized with ketamine/xylazine, and intranasal inoculation was performed by the administration of 20 µL (2×10^6^ CFU/dose) of the inoculum, applying 10 µL into each nare. Mice were monitored for 6 days, and moribund mice were euthanized (30).

### Opsonophagocytic Killing Assays

Opsonophagocytic killing (OPK) assays were performed following methods as previously described (42). While OPK activity below 50% can be statistically significant, OPK activity is generally only biologically significant when above 50% (24).

### Splenocyte isolation and co-culture assays for IL-17 secretion

Splenocytes were isolated and stimulated with vaccine antigens as previously described (23). Briefly, spleens were aseptically removed and suspended in PBS containing 2% heat-inactivated FCS. Spleens were disaggregated by passing through 100-micron nylon screens into a petri dish. Erythrocytes were lysed using a Mouse Erythrocyte Lyse Kit (R&D Systems) per the manufacturer’s protocol. Cells were centrifuged and resuspended in 5 mL cR10 (RPMI with 2 mM glutamine, 1 mM sodium pyruvate, 1X non-essential amino acids solution, 1x penicillin/steptomycin, 55 µM 2-mercaptoethanol, and 10% heat-inactivated FCS, all from Invitrogen), adjusted 1×10^6^ cells/mL, and seeded into 96-well round-bottom polystyrene plates. Cells were stimulated with 1 ug/mL protein for 7 days at 37°C in 5% CO_2_. Supernatants were collected and assayed for IL-17 by ELISA (R&D Systems).

### ELISAs

ELISAs to measure PopB-, OprF/I-, or whole *P. aeruginosa-*specific IgG titers were performed with plates (Immulon 4HBX) coated with proteins at 1 _μ_g/mL or live bacteria, as previously described (43). *P. aeruginosa* strain N13 was grown in LB containing 5 mM EGTA to induce the type III secretion system (TTSS) to maximize PopB expression (44).

### OprF/I-IFN-γ binding inhibition assays

Assays for the inhibition of binding of human IFN-γ to plate-bound OprF/I was performed as previously reported (19). Briefly, 100 µL of OprF/I (1 _μ_g/mL) was applied to the wells of an ELISA plate (Immulon 4HBX) and incubated overnight at 4 C. After washing with PBST, the plates were blocked with 5% BSA in PBS for 1 h at 37 C. OprF/I-coated plates were then incubated with 100 µL of 1:2 dilutions of antisera for 2 h at 37 C. After washing, 100 µL of human IFN-γ (0.5 _μ_g/mL) (R & D Systems) was added to the wells and incubated overnight at 4 C. The bound IFN-γ was detected using the detection antibody and substrate system using human IFN-γ DuoSet reagents (R & D Systems) according to the manufacturer’s protocol.

### Statistical analyses

All analyses were performed using Prism (Graphpad Software). Survival data were analyzed with the Kaplan-Meier method and log-rank tests. Parametric data were analyzed by t test or by ANOVA with Dunnett’s *post-hoc* multiple comparison test. Nonparametric data were analyzed by Mann Whitney U test or Kruskal Wallis test with Dunn’s multiple comparison test. EC_50_ was determined by nonlinear regression with 95% confidence intervals based on profile likelihood. We determined EC_50_ values to be statistically significant if the 95% CIs did not overlap.

## Acknowledgments

This work was supported in part by the US Department of Defense in an Investigator-Initiated Award of the Peer Reviewed Medical Research Program, grant no. PR181874, W81XWH-19-1-0208 (to G.P.P.), by the Technology Development Fund of the Technology and Innovation Development Office (TIDO) at Boston Children’s Hospital (to G.P.P.), by the Richard A. and Susan F. Smith President’s Innovation Award (to G.P.P.), and by the Translational Research for Infection Prevention in Pediatric Anesthesia and Critical Care (TRIPPACC) Program of the Department of Anesthesiology, Critical Care and Pain Medicine at Boston Children’s Hospital (to G.P.P.).

## Figure Legends

**Figure S1.**
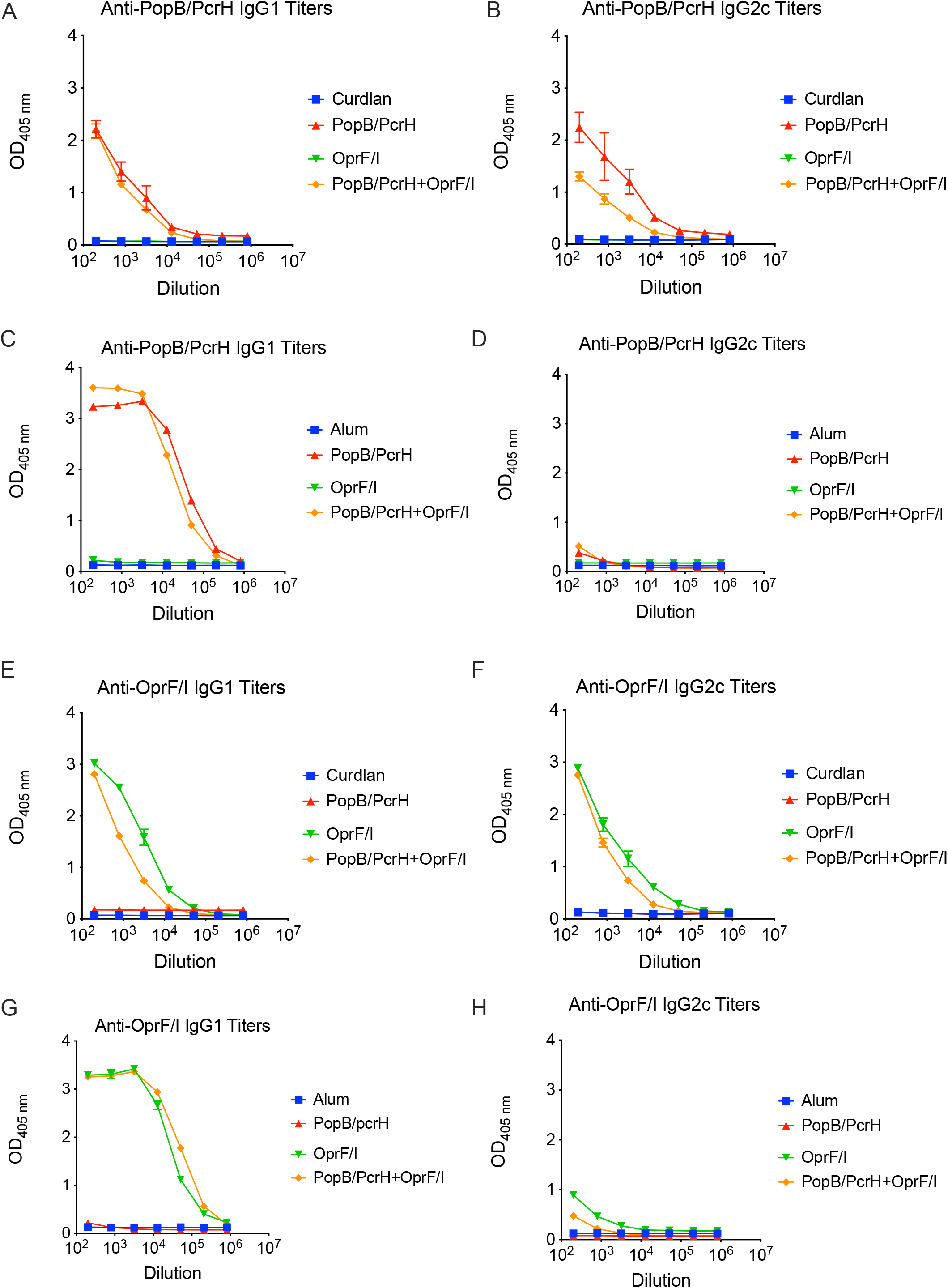
Vaccination with PopB/PcrH, OprF/I, or both, generates antigen-specific IgG1 and IgG2c responses. Mice were immunized either intranasally (A,B,E,F) or subcutaneously (C,D,G,H) with adjuvant alone (curdlan or Alum), adjuvant + 30 µg PopB/PcrH, adjuvant + 30 µg OprF/I, or adjuvant + 30 µg PopB/PcrH + 30 µg OprF/I, and sera were collected three weeks after the last immunization. Anti-PopB/PcrH (A,C) and anti-OprF/I (E,G) IgG1 titers and anti-PopB/PcrH (B,D) and anti-OprF/I (F,H) IgG2c titers were measured using ELISA. Sera from 3-4 mice per group were pooled and measured in duplicate, and means are plotted with SD as error bars (error bars are smaller than symbol at many points). Data are representative of at least two independent experiments.

**Figure S2.**
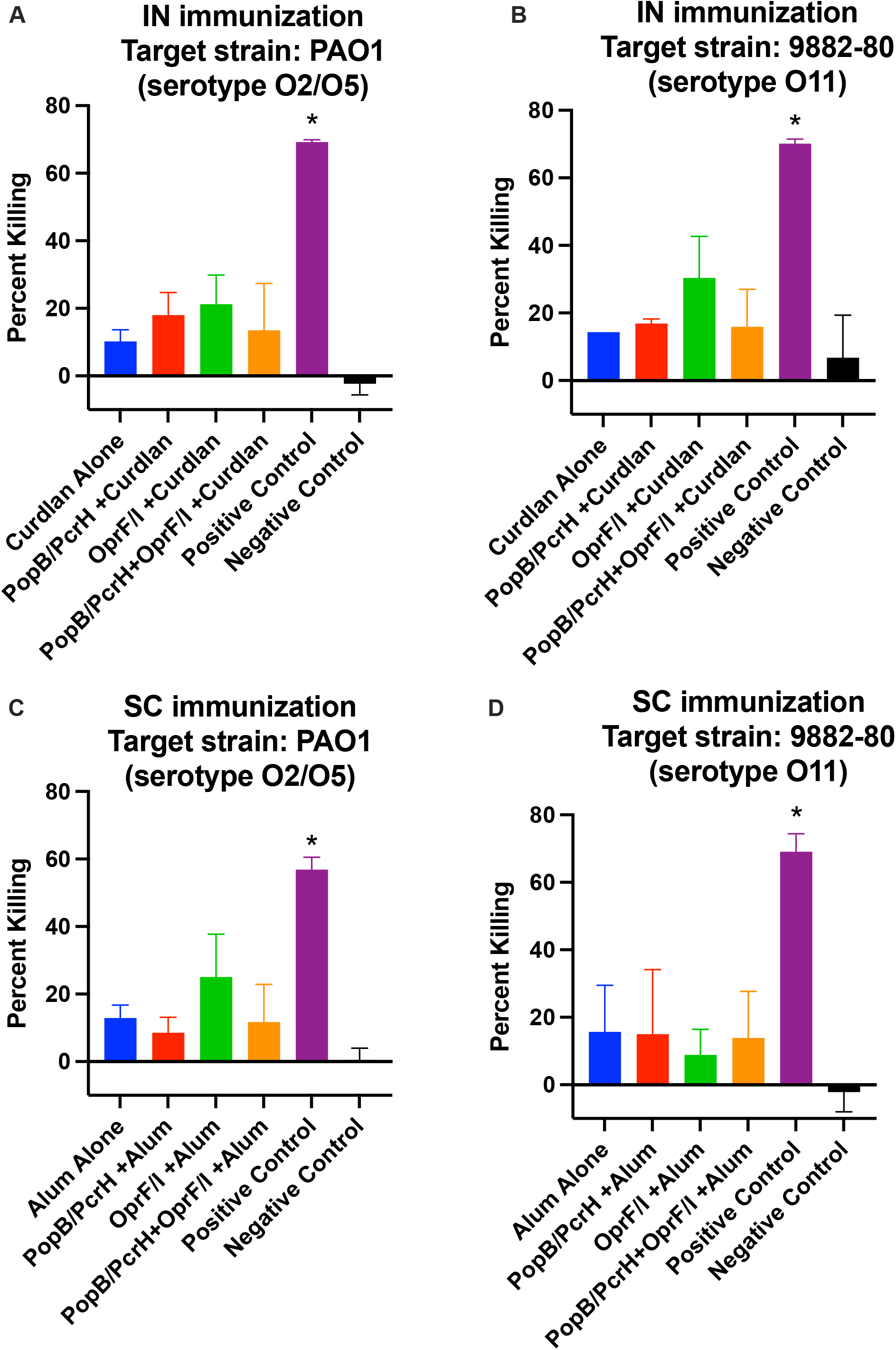
Vaccination with PopB/PcrH, OprF/I, or both, do not generate significant opsonophagocytic killing activity. Mice were immunized either intranasally (A,B) or subcutaneously (C,D) with adjuvant alone (curdlan or Alum), adjuvant + 30 µg PopB/PcrH, adjuvant + 30 µg OprF/I, or adjuvant + 30 µg PopB/PcrH + 30 µg OprF/I, and sera were collected three weeks after the last immunization. *P aeruginosa* strain PAO1 (serotype O2/O5) and strain 9882-80 (serotype O11) were target strains, and sera were used at 1:30 dilution. As a positive control, an anti-Psl monoclonal antibody Cam003 (10µg/ml) was used. R347, human monoclonal antibody to HIV gp120 (10µg/ml) was used as a negative control. *denotes *p*<0.05 by one-way ANOVA followed by Dunnett’s *post-hoc* multiple comparison test when compared to the adjuvant only vaccine group (curdlan or Alum).

**Table S1.** EC50 calculations for titers measured by ELISA.

**A. Total IgG:**
|  | Vaccine Groups | Anti-PopB/PcrH IgG |  | Anti-OprF/I IgG |  | Anti- *P. aeruginosa* IgG |  |
| --- | --- | --- | --- | --- | --- | --- | --- |
|  |  | EC<sub>50</sub> | 95% CI | EC<sub>50</sub> | 95% CI | EC<sub>50</sub> | 95% CI |
| SC | Alum | NC | NC | NC | NC | NC | NC |
|  | PopB/PcrH | 33,683 | 29,877 - 38,103 | NC | NC | 10 | 4 - 17 |
|  | OprF/I | NC | NC | 9,330 | 7,760 - 11,133 | 1,832 | 1,667 - 2,019 |
|  | PopB/PcrH + OprF/I | 3,007 | 2,718 - 3,313 | 15,699 | 14,107 - 17,549 | 2,042 | 1,911 - 2,184 |
| IN | Curdlan | NC | NC | NC | NC | NC | NC |
|  | PopB/PcrH | 3,527 | 3,340 - 3,721 | NC | NC | 94 | 82 - 107 |
|  | OprF/I | NC | NC | 7,992 | 7,616 - 8,381 | 72 | 61 - 83 |
|  | PopB/PcrH + OprF/I | 3,499 | 3,244 - 3,770 | 8,856 | 8,326 - 9,407 | 190 | 163 - 220 |

**B. IgG Subclasses:**
|  | Vaccine Groups | Anti-PopB/PcrH IgG1 |  | Anti-OprF/I IgG1 |  | Anti-PopB/PcrH IgG2c |  | Anti-OprF/I IgG2c |  |
| --- | --- | --- | --- | --- | --- | --- | --- | --- | --- |
|  |  | EC<sub>50</sub> | 95% CI | EC<sub>50</sub> | 95% CI | EC<sub>50</sub> | 95% CI | EC<sub>50</sub> | 95% CI |
| SC | Alum | NC | NC | NC | NC | NC | NC | NC | NC |
|  | PopB/PcrH | 37,957 | 34,291 - 42,045 | NC | NC | 274 | 13 - 572 | NC | NC |
|  | OprF/I | NC | NC | 28,524 | 24,859 - 32,700 | ND | NC | 113 | NC |
|  | PopB/PcrH + OprF/I | 18,829 | 16,762 - 21,227 | 53,868 | 48,270 - 60,591 | 3.3 | NC | 0.4 | NC |
| IN | Curdlan | NC | NC | NC | NC | NC | NC | NC | NC |
|  | PopB/PcrH | 526 | NC | NC | NC | 1,974 | NC | NC | NC |
|  | OprF/I | NC | NC | 3,013 | 2,593 - 3,449 | NC | NC | 96 | NC |
|  | PopB/PcrH + OprF/I | 69 | NC | 472 | 312 - 623 | 722 | 49 - 1,429 | 192 | 13 - 443 |

Abbreviations:
NC= not calculated; EC<sub>50</sub>= half maximal effective concentration; 95% CI = 95% of confidence interval

